# TALE and Hox Transcription Factors Control Adult Behaviors in Zebrafish

**DOI:** 10.64898/2026.03.16.712211

**Authors:** Austin Adkins, Katelyn Glowinski, Yong-Il Kim, Ethan Wright, Cameron E Bennett, Jessica C Nelson, Charles G Sagerström

**Affiliations:** Section of Developmental Biology, Department of Pediatrics, University of Colorado Anschutz, Aurora, CO, USA; Department of Cell and Developmental Biology, University of Colorado Anschutz, Aurora, CO, USA

**Keywords:** Mental Health, TALE and Hox genes, Transcriptional Regulation, Gene Function in Behavior

## Abstract

Behavioral dysfunction is a common characteristic of many mental health disorders. While the causes of these disorders vary, aberrant behaviors may arise from alterations in transcriptional regulation already during early neural development. Because transcription factors (TFs) often belong to families of closely related members, disruption of a single TF may indirectly influence the functionality of other family members. Consequently, mutations in TFs within the same family may lead to overlapping, yet distinct, phenotypes. This feature of TF function has important implications for understanding behavioral phenotypes, but detailed analyses across a single TF family are still lacking. In this study, we present a comprehensive behavioral analysis of adult zebrafish harboring mutations in individual members of the TALE and Hox TF families, that are essential for nervous system development. Using a battery of validated behavioral assays, we uncover elevated stress responses among all TF mutant lines, as well as TF-specific dysregulation in coping strategies, social interaction, learning/memory, and endurance and locomotion. The shared behavioral abnormalities across mutants suggests TF family members converge on core developmental pathways for stress-related behavioral regulation, while mutation-specific phenotypes indicate unique roles for individual TFs in fine-tuning neural function. Our findings provide a systematic behavioral characterization of TALE and Hox mutants in a vertebrate model and provide a framework for understanding how genetic variation within TF families may differentially contribute to vulnerability for mental health disorders.

## 1. Introduction

Behavioral dysfunction is a hallmark of many mental health disorders, often manifesting as impairments in cognition, emotional regulation, and social interaction. Aspects of these altered behaviors may stem from disruptions to early neural development. During embryogenesis, the formation of functional neuronal circuits depends on the spatiotemporal coordination of gene expression that governs neuronal specification, migration, synaptogenesis, and circuit refinement (Lupo et al. 2006; Goodman and Bonni 2019; Pei et al. 2021). Transcription factors (TFs) are central to these developmental programs, regulating diverse gene networks that orchestrate patterning and cell fate decisions (Arendt et al. 2016; Lambert et al. 2018). Disruptions in TF function, or their epigenetic regulation, have been implicated in both syndromic and idiopathic mental health and neurodevelopmental disorders, including those involving chromatin remodelers (e.g., CHD8 (Shi et al. 2023), MECP2 (Amir et al. 1999)) and neural TFs such as TBR1 (Huang and Hsueh 2015) and Pax5 (Gofin et al. 2022). Importantly, many TFs belong to conserved families, where individual members bind similar DNA motifs and exhibit overlapping but not identical functions.

Consequently, loss of one TF may lead to family members with similar binding specificity taking its place at regulatory elements. While this may partially compensate for the inactive TF, even subtle functional differences between family members could lead to altered gene expression programs and disrupted neural development. Further, TFs frequently act together in multiprotein complexes, meaning that loss of one TF may not simply inactivate the complex, but alter its function. Such altered complexes may locate to inappropriate regulatory elements and/or display aberrant activating or repressive functions. These features of TFs suggest that the phenotype resulting from mutations in one TF is not due simply to the loss of that TF, but possibly also to aberrant activities of related TFs and complexes. Despite such atypical TF activities potentially introducing new layers of dysregulation during development (Spitz and Furlong 2012), and having important implications for our understanding of the genetic basis for disordered human behaviors, we do not know if disruption of closely related TFs results in behavioral phenotypes with identical, or partially overlapping, phenotypes.

Members of the TALE (<u>T</u>hree <u>A</u>mino acid <u>L</u>oop <u>E</u>xtension) homeodomain TF family—Meis, Pbx, Prep, and TGIF—are key regulators of neurodevelopment, with well-established roles in neural patterning, neuronal progenitor differentiation, and axonal pathfinding (Waskiewicz et al. 2001, 2002; Choe et al. 2002; Gongal and Waskiewicz 2008; Ladam et al. 2018), as well as in neurogenesis, cell proliferation, neuronal circuit formation, and synaptic plasticity (Agoston et al. 2014; Grebbin et al. 2016; Hau et al. 2017, 2021; Ladam et al. 2018; Müller et al. 2024; Kittke et al. 2024). These TFs frequently act in complexes with Hox proteins, modulating their binding specificity and transcriptional output (Mann and Affolter 1998; Bobola and Sagerström 2024). As such, Pbx and Meis, as well as Hox TFs, are recognized as being implicated in mental health disorders (Coe et al. 2019; Fu et al. 2022; NIMH 2023) and there is also strong evidence for the involvement of Prep (Sanchez-Font 2003; Deflorian et al. 2004) and TGIF (Gripp et al. 2000; Knepper et al. 2006) TFs in neurodevelopmental and mental health disorders. Because the TALE family includes closely related TFs, and because TALE TFs form complexes with each other (both Meis and Prep form complexes with Pbx (Chang et al. 1997; Berthelsen et al. 1999)) as well as with Hox TFs (Ferretti et al. 2005; Merabet and Mann 2016), the TALE and Hox families represent a useful model to determine if closely related TF family members have similar or distinct roles in controlling behaviors relevant to mental health.

Despite extensive studies on their neurodevelopmental roles, little is known about how disruptions in TALE and Hox TF function affect behavioral outcomes. Past studies in mice showed that a Meis1 mutation results in altered motor phenotypes (Salminen et al. 2017; Kittke et al. 2024) and nociceptor development (Cao et al. 2022), indicating a role for this TF in neural circuits underlying physiological processes and complex behaviors. Here, we extend this prior work and provide a comprehensive examination of the roles of the remaining TALE and several Hox TFs in establishing adult behaviors. We find that adult TALE and Hox mutant zebrafish display persistent behavioral abnormalities, including altered stress responses, impaired memory processing, and dysregulation of social patterning. TALE and Hox mutants show some overlapping presentations of these behavioral alterations, while other phenotypes are unique to specific TF family members. Notably, these phenotypes are not accompanied by obvious morphological defects, suggesting subtle but functionally significant circuit-level disruptions. To our knowledge, our work represents the first comprehensive characterization of adult behavioral phenotypes in zebrafish mutants for members of the TALE TF family and two early acting Hox TFs.

## 2. Materials and Methods

### 2.1 Subjects

Aged-matched adult 3-9 month old AB strain wildtype (WT) and mutant zebrafish lines (maintained on the AB background) were bred and raised in our zebrafish housing facility. We have previously reported the *hoxb1a^um197^* and *hoxb1b^um191^* mutant lines (Weicksel et al. 2014), and we obtained the *TGIF^fh258^* mutant line (an ENU-induced early STOP codon; (Lenkowski et al. 2013)) from the Zebrafish International Resource Center (ZIRC). We generated the *pbx2^co126^* and *pbx4^co127^* mutant lines using CRISPR-induced indels that produce premature STOP codons, using methods we reported previously (Ghosh et al. 2018; Maurer and Sagerström 2018; Yildiz et al. 2019). All mutations represent putative null alleles as they introduce STOP codons upstream of the DNA-binding homeodomain (**Supplemental Table 1**). Fish were fed twice per day (morning and evening) with GEMMA Micro 300 (Skretting; Westbrook, ME). The housing and behavioral testing facilities were kept on a 14:10 light:dark cycle and 30-40% ambient humidity, and water temperatures maintained at 28°C ± 0.5°C. Fish used for behavioral testing were not subjected to breeding crosses and experienced minimal handling, limited to brief transfers during tank relocation prior to behavioral assessment. All procedures were conducted in accordance with the National Institutes of Health Guide for the Care and Use of Experimental Animals and were approved by the University of Colorado’s Institutional Animal Care and Use Committee (Protocol#: 870).

### 2.2 Behavioral Testing

Fish designated for individual testing were separated (1:1 ratio Male:Female, n= 24 total) and singly housed 48hr prior to the first day of behavioral assessment. For group testing, fish were relocated into groups of 6 (1:1 ratio Male:Female, n= 24 total) and allowed to acclimate one week prior to behavioral assessment. For all behavioral paradigms on testing days, all fish were allowed to acclimate to the behavioral testing room for 2hr prior to testing. All testing occurred at the start of the 3^rd^ hr of lights on. Video recording occurred for the duration of testing for each paradigm and were analyzed using the Noldus EthovisionXT® software (Version 17.5.1718; Leesburg, VA). Fish were transported in their home tanks and transferred to each testing chamber via the pour off method to minimize stress caused by handling and transportation (Shishis et al. 2023). An experimental timeline is detailed in **Supplemental Fig. 1**. Details for each behavioral task are outlined below.

#### 2.2.1 Novel Object Recognition (NOR)

Individual adult zebrafish were placed into a custom 28cm x 28cm x 15cm acrylic arena filled to a water depth of 6cm and allowed to explore for 10min. On Day 1 the chamber contained two objects (0.5in flat bottomed glass marbles, Amazon, ASIN: B0986V6QTG) of identical color placed 10cm from the top left (Object 1) and bottom right (Object 2) corners. On Day 2, Object 2 was replaced with a different colored glass marble (i.e., the novel object). The fish were again allowed to explore for 10min. Time spent exploring the novel object versus the familiar one is characterized as an index of memory (May et al. 2016; Faillace et al. 2017; Gaspary et al. 2018). This experiment was performed either with the two objects being blue on Day 1 and Object 2 switched to green on Day 2, or with the two objects being red on Day 1 and Object 2 switched to green on Day 2. The time spent per visit with each object on each day were measured using the “In Zone” feature in EthovisionXT. To calculate the difference in time spent with each object on Day 2 versus Day 1, we subtracted the average time spent per visit to the zone surrounding each object on Day 1 from the average time spent per visit to the zone surrounding that object on Day 2. If the difference is a positive number, more time was spent per visit to the zone surrounding that object on Day 2; if it is a negative number, less time was spent per visit to the zone surrounding that object on Day 2 (**Fig. 1a**).

**Fig. 1.**
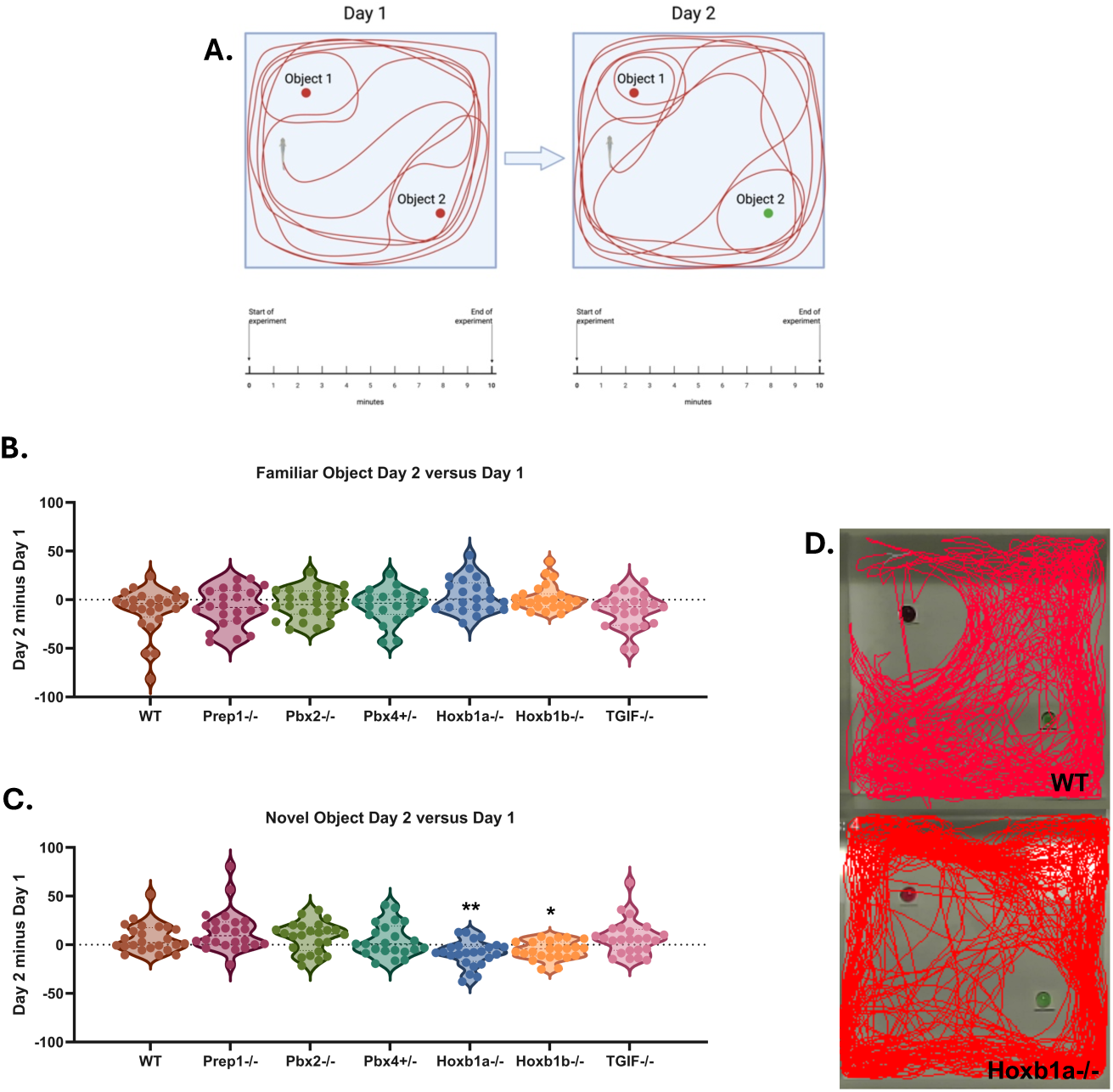
Object Recognition Memory is Negatively Affected by Hox Mutations **a)** Schematic of the novel object recognition (NOR) paradigm. Violin graphs showing the time spent around **b**) the familiar object (Object 1) and **c**) the novel object (Object 2) on the testing day (Day 2) relative to the training day (Day 1). Each data point represents one animal. For these plots, time spent around an object on Day 1 was subtracted from the time spent around that object on Day 2. If the difference is a positive number, more time was spent around the object on Day 2; if it is a negative number, less time was spent around the object on Day 2. **d)** Representative line traces of individual fish in the NOR paradigm showing differences in WT versus Hoxb1b-/- lines on Day 2. Data were analyzed using one-way ANOVAs. Significant differences compared to WT: *p<0.05, **p<0.01.

#### 2.2.2 Open Field (OF)

Individual adult zebrafish were placed into a custom 28cm x 28cm x 15cm acrylic arena filled to a water depth of 6cm and allowed to explore for 10min. More time spent around the perimeter of the tank (thigmotaxis (Prut and Belzung 2003; Champagne et al. 2010)) was considered as increased anxiety-like behavior compared to time spent in the center of the tank. Swim paths of each fish, percent of time spent in a freezing or hyperactive state (using the “Activity State” feature in EthovisionXT), and time spent in different areas of the tank (using the “In Zone” feature in EthovisionXT) were measured (**Fig. 2a**).

**Fig. 2.**
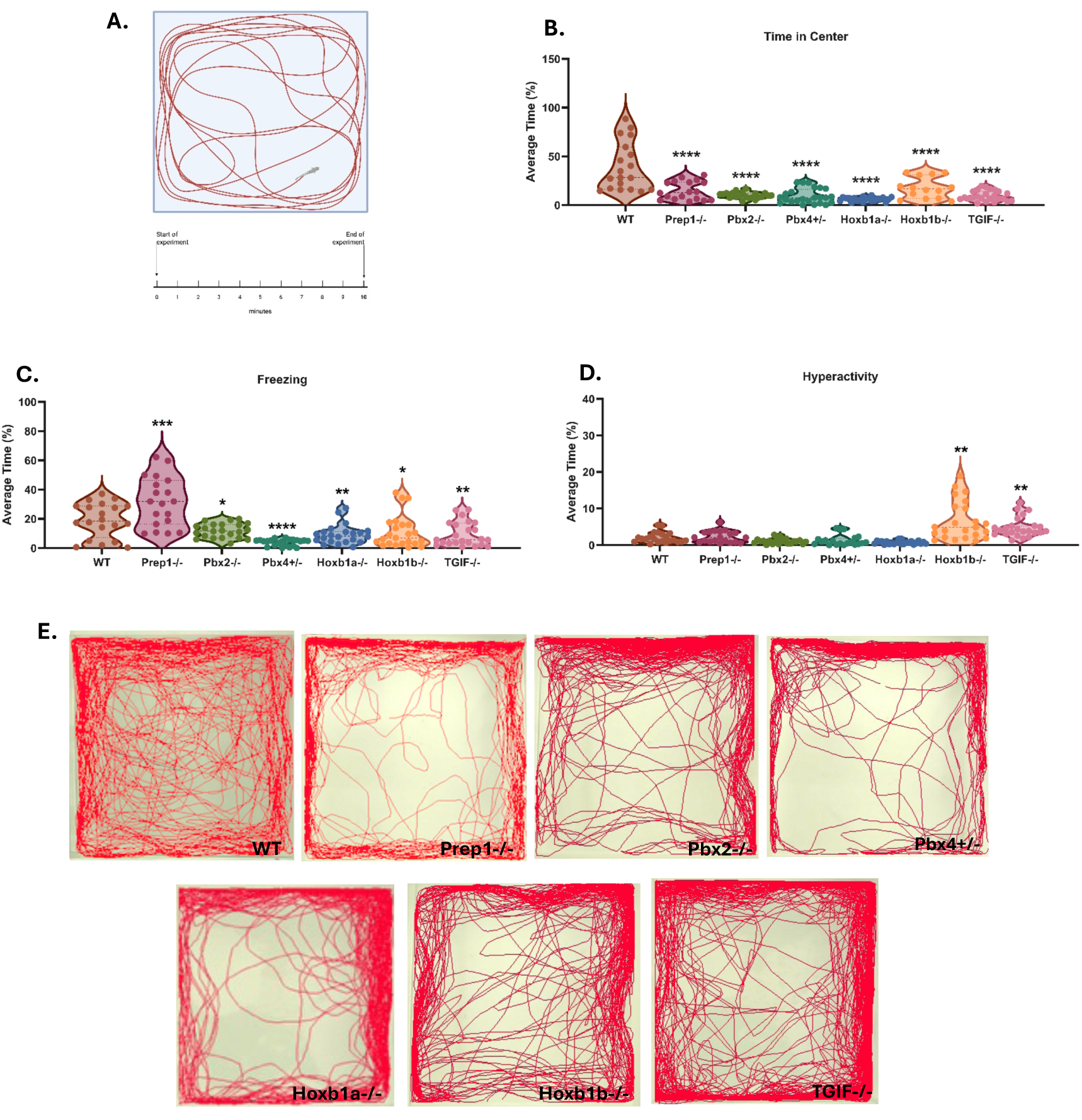
Stress-Responsive Behaviors in the Open Field Vary Across TALE and Hox Mutant Lines Despite Shared Thigmotaxis **a)** Schematic of the open field paradigm. Violin graphs showing the **b)** percent time, averaged across the 10min time period, spent in the center of the open field, **c)** percent time, averaged across the 10min time period, spent freezing, and **d)** percent time, averaged across the 10min time period, spent hyperactive. Each data point represents one animal. **e)** Representative line traces of individual fish in the OF paradigm across genotypes, highlighting differences in spatial exploration. Data were analyzed using one-way ANOVAs. Significant differences compared to WT: **p<0.01, ***p<0.001, and ****p<0.0001

#### 2.2.3 Novel Tank Dive (NTD)

Individual zebrafish were placed into a custom 5cm x 20cm x 15cm arena filled to a water depth of 13cm.

Fish were allowed to explore the arena for 10min. Typically, when introduced to a vertical novel environment, zebrafish will naturally dive to the bottom of the tank, then gradually increase their vertical swimming and exploration over time. This gradual increase is interpreted as a reduction in anxiety (Levin et al. 2007; Cachat et al. 2010; Gebauer et al. 2011). As such, due to the vertical distribution of the testing chamber (as opposed to horizontal distribution in OF), more time spent in the bottom of the tank is indicative of increased thigmotaxis. Decreased time spent in the lower portion of the tank is considered indicative of reduced anxiety. The duration of time spent in defined zones (upper, middle, and lower third of the tank), the number of zone transitions during the monitoring period (using the “In Zone” feature in EthovisionXT), and percent of time spent in a freezing or hyperactive state (using the “Activity State” feature in EthovisionXT) were measured (**Fig. 3a**).

**Fig. 3.**
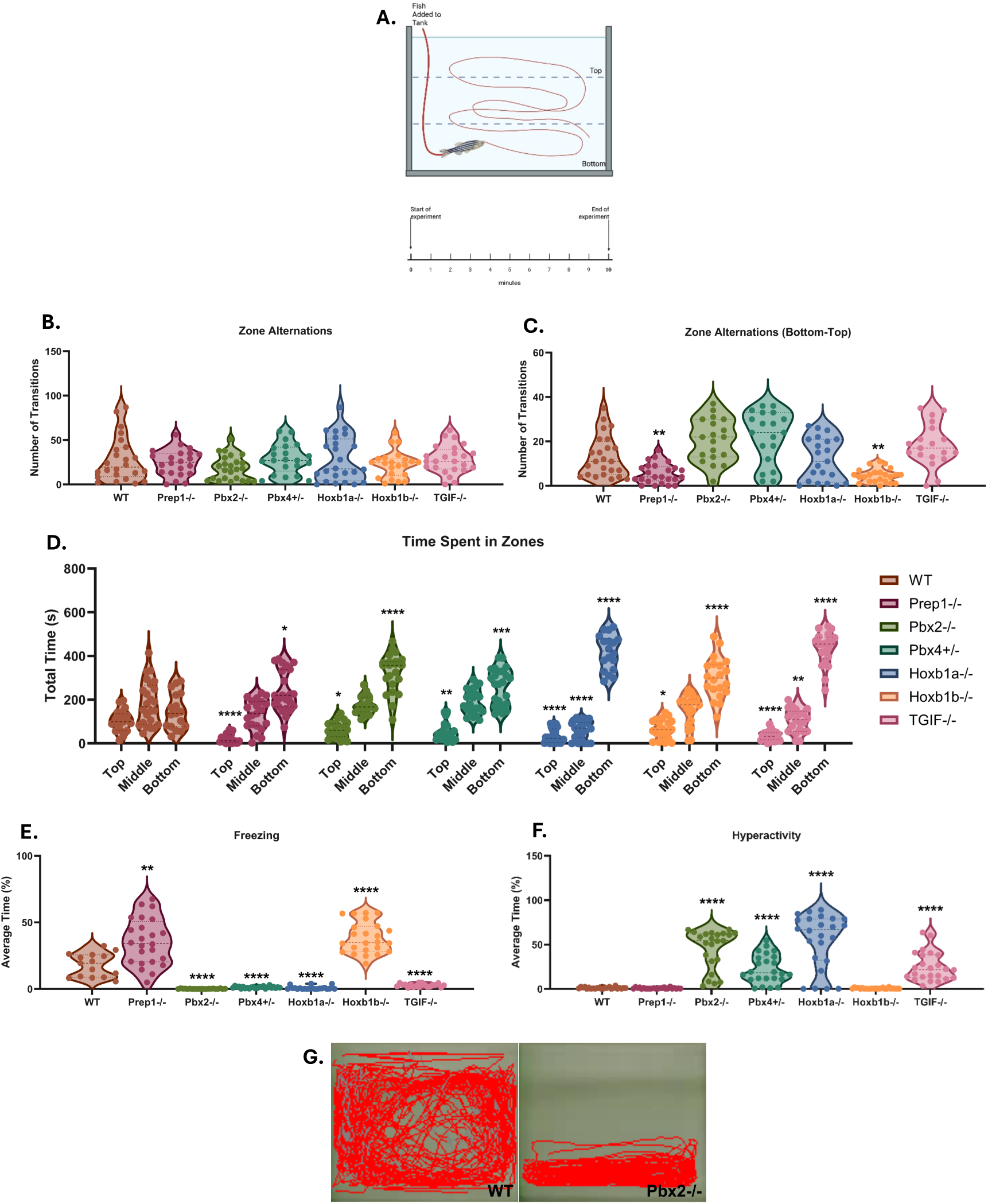
Stress-Responsive Behaviors in the Novel Tank Dive Vary Across TALE and Hox Mutant Lines **a)** Schematic of the novel tank dive paradigm. Violin graphs showing **b)** overall number of zone alternations, **c)** number of transitions from the bottom to the top of the water column, **d)** total time spent in each zone of the water column, **e)** percent time, averaged across the 10min time period, spent freezing, and **f)** percent time, averaged across the 10min time period, spent hyperactive. Each data point represents one animal. **g)** Representative line traces of individual fish in the NTD paradigm showing differences in WT versus Pbx2-/-. Data were analyzed using one-way ANOVAs (**b**-**c**, **e**-**f**) or two-way ANOVA (**d**). Significant differences compared to WT: *p<0.05, **p<0.01, ***p<0.001, and ****p<0.0001

#### 2.2.4 Sociability

To assess social behavior, we examined adult zebrafish under baseline conditions and during a net chase paradigm (**Fig. 4a**). Shoaling is a prominent, innate social behavior in zebrafish in which individuals aggregate, maintain consistent spacing, and respond to each other’s movements without necessarily swimming in unison (Pitcher and Parrish 1993; Miller and Gerlai 2007). Shoaling patterns are sensitive to environmental and social context as groups often tighten under threat, while reduced shoal cohesion can indicate decreased anxiety-like behavior (Speedie and Gerlai 2008). To measure shoaling behavior, fish were video recorded in groups of 6 adults (1:1 ratio of males and females) for 10min at baseline in their 3L home tank (Aquaneering; San Marcos, CA). Inter-individual distance (IID) was measured using the “Distance Between Subjects” feature in EthovisionXT to assess sociability within the group, with a decreased IID as a measure of increased sociability/group cohesion and increases in IID as a measure of lack of group cohesion, and therefore potential deficits in social behaviors (Buske and Gerlai 2012). In addition to IID, spatial distribution within the tank was also analyzed using the “In Zone” feature in EthovisionXT. Shoals that disperse evenly throughout the tank are considered to reflect a more relaxed, low-stress state. In contrast, shoals that cluster tightly in one region of the tank may indicate heightened stress or anxiety within the group (Speedie and Gerlai 2008).

**Fig. 4.**
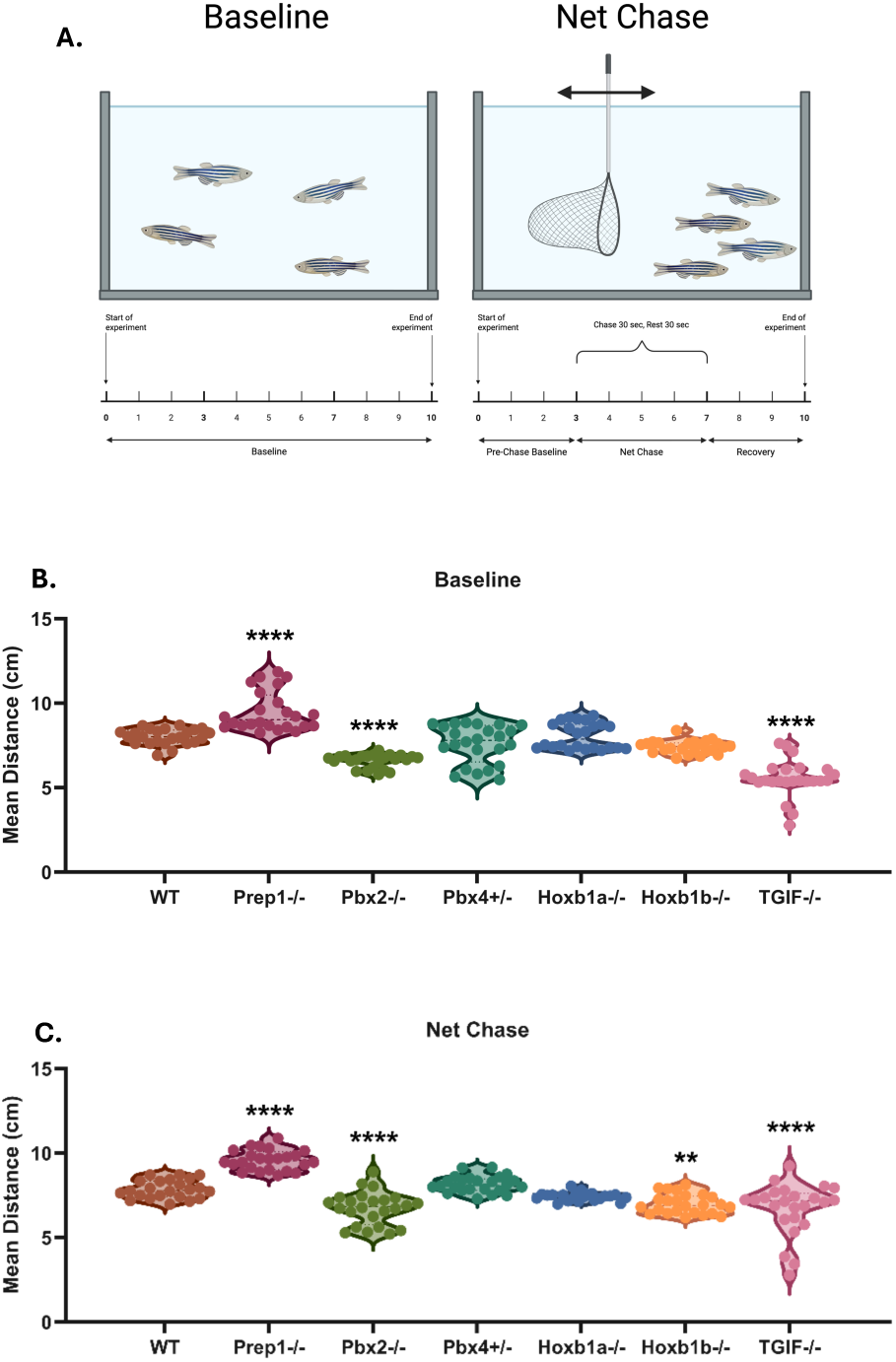
Sociability Differs at Baseline and Net Chase Between TALE and Hox Mutants **a)** Schematic of the sociability paradigms. Violin plots showing **b)** the average inter-individual distance (IID) between individuals in a group during baseline, **c)** the average inter-individual distance (IID) between individuals in a group during net chase. For each animal, we calculated the average IID between that animal and the other five animals in the tank. Each data point reflects one animal’s value. Data were analyzed using one-way ANOVAs. Significant differences compared to WT: **p<0.01, ****p<0.0001

To asess the effect of stress on social behavior, we utilized the common net chasing paradigm which mimics the fish being pursued and captured using a net (Ramsay et al. 2009; Aponte and Petrunich-Rutherford 2019). In our modified version of this paradigm, following a 3min pre-net chase period, fish are chased with a net that is swept across the tank, captured, and pulled out of the water for a brief period (<5s) and immediately freed from the net and returned to the tank. Net chasing occurred at the start of the 3^rd^ min and ended after the 7^th^ min for a total of 5 sequential net chases. Net chasing occurred repeatedly for a 30s period, followed by a 30s rest period, each minute. After the last net chase, the fish received a 3min post-net chase period to measure recovery and a return to homeostasis. Groups of fish were video recorded for 10min and behavior was measured as described above. Because the net chase paradigm also causes the fish to expend energy to avoid being captured, additional measures were used to distinguish physiological responses in each line, including the number of lethargic individuals, overall activity, and latency to return to homeostasis (indicated by returning to normal shoaling behavior and moving up the water column) using the “Activity State” feature in EthovisionXT.

### 2.3 Statistical Analyses

Data were analyzed with one-way or two-way ANOVAs (depending on design), followed by Tukey’s *post hoc* comparisons test (when indicated by a significant ANOVA) to test for TF mutant-specific effects on individual behavioral metrics compared to WT. All ANOVAs were generated using GraphPad PRISM software (Version 9.4.1).

## 3. Results

Mental health disorders frequently involve disruptions across multiple behaviors, including memory and cognition, stress responsivity, and social interaction (Chand and Marwaha 2025; Hany and Rizvi 2025; Hodis et al. 2025; Musullulu 2025). These are often interrelated. For instance, impairments in recognition memory and cognitive flexibility are core features of disorders such as schizophrenia and autism spectrum disorder (ASD) (Oh et al. 2017; Desaunay et al. 2023), while altered responses to acute stress—ranging from passive freezing (Schmidt et al. 2008; Volchan et al. 2017) to hyperarousal (Cantor 2009; Mahoney et al. 2018)—are characteristic of anxiety, depression, and PTSD. Furthermore, deficits or dysregulation in social engagement are hallmark symptoms across a wide range of neuropsychiatric conditions, including ASD and affective disorders (Yamada et al. 2023; Fulford and Holt 2023). This interrelatedness has been predicted to reflect underlying vulnerabilities in neuronal circuit development and function, which require proper control of gene expression programs. Accordingly, TFs have been implicated in various mental health disorders (Coe et al. 2019; Fu et al. 2022; NIMH 2023). TALE and Hox TFs are central regulators of brain development, orchestrating neuronal specification, circuit formation, and connectivity. These TFs have also been implicated by the NIMH as associated with mental health disorders (NIMH 2023), yet their roles in the mechanisms linking TF function to behavior remain largely unexplored. In particular, it is unclear how closely related TFs—sharing sequence homology, DNA recognition motifs, and cofactors—may produce overlapping or distinct effects on behavioral outcomes. To address this gap, we systematically examined adult behaviors in zebrafish mutants for TALE and Hox TFs.

The TALE TF family consists of four members – Pbx, Prep, Meis and TGIF (Bobola and Sagerström 2024). While there are paralogous genes for each TALE member – e.g., there are four Pbx paralogs in zebrafish – this study focused on examining the predominant paralog (based on gene expression and severity of loss-of-function phenotype (Waskiewicz et al. 2001, 2002; Deflorian et al. 2004; Gongal and Waskiewicz 2008; Lenkowski et al. 2013) for each TF (i.e., Pbx4, Prep1, and TGIF1). Since the Pbx4 null mutant is not homozygous viable, we also examined the next most common paralog, Pbx2 (Waskiewicz et al. 2002), which is homozygous viable. Since Meis genes have been examined previously(Salminen et al. 2017; Delgado et al. 2021; Kittke et al. 2024), we omitted them from our analysis. Because TALE TFs form complexes with early-acting Hox TFs (Bobola and Sagerström 2024), we also examined mutants for the zebrafish Hoxb1a and Hoxb1b TFs. To probe the behaviors of these mutants, we employed a battery of ethologically grounded assays in zebrafish that collectively model recognition memory (novel object paradigm), stress responsivity (open field, novel tank dive, and net chase tasks), and social cohesion (baseline and net chase paradigms). These tasks enable the quantification of spontaneous and elicited behaviors, allowing us to examine both baseline functioning and responses to environmental perturbation.

### 3.1 Object Recognition Memory is Affected by Hoxb1a and Hoxb1b, but not TALE, Mutations

Novel object recognition (NOR) is a simple task used to quantify memory function in zebrafish, comparable to task performance in rodents (May et al. 2016; Gaspary et al. 2018). For this task, zebrafish were allowed to familiarize themselves with two identical objects on Day 1, and we assessed their response when one object was replaced with a novel one on Day 2 (**Fig. 1a**, see Methods for details). When analyzing the difference in time spent with objects between days, we find that WT fish spend more time with the novel object than the familiar one on Day 2 (**Fig. 1b-d**), suggesting intact recognition memory and a natural exploratory drive toward the novel stimulus. Both Hoxb1a and Hoxb1b mutant fish spent significantly less time around the novel object compared to WT fish on Day 2 (p<0.01 and p<0.05, respectively; **Fig. 1c-d**). This observation indicates that the Hoxb1a and Hoxb1b genes may be required for normal recognition memory, but we cannot fully exclude other possible defects, such as effects of innate color preferences (Faillace et al. 2017). In contrast, Prep1, Pbx2, Pbx4 and TGIF1 mutants do not show recognition memory phenotypes, suggesting that the observed deficits are specific to Hox disruption rather than a general property of TALE and Hox mutants.

### 3.2 Stress-Responsive Behavioral Profiles and Coping Strategies Vary Across TALE and Hox Mutant Lines

To assess stress-related behavior, we analyzed performance across both the open field (OF) and novel tank dive (NTD) tasks. In the OF task (**Fig. 2a**, see Methods for details), WT fish explored the arena broadly, spending time in both the center and the periphery (**Fig. 2b, e**), a pattern consistent with a low-stress state and robust exploratory drive. Contrastingly, all TALE and Hox mutant lines exhibited a significant increase in thigmotactic behavior, spending less time in the center than the periphery of the arena (**Fig. 2b, e**) compared to WT controls (p<0.0001 for all genotypes), indicative of increased anxiety-like behavior (Prut and Belzung 2003; Champagne et al. 2010).

Despite this shared elevation in anxiety-like behavior, the nature of the stress response differed across mutant lines. Prep1 mutants showed a significant increase in freezing behavior relative to WT (p<0.001), consistent with a passive coping strategy or heightened fear response (Ziv et al. 2013). All other mutant lines showed a significant decrease in freezing (**Fig. 2c**). In the TGIF1 (p<0.01) and Hoxb1b (p<0.001) mutants, the decreased freezing behavior was paired with significantly increased hyperactivity (**Fig. 2d**), suggestive of an active or hyper-aroused response to stress (Sumathipala et al. 2024). In the other lines (Pbx2, Pbx4, and Hoxb1a) the decrease in freezing was not accompanied by increased activity levels (**Fig. 2c-d**), indicating a subtler alteration in stress reactivity.

Similar to OF, the NTD task uses a vertical distribution in a novel environment (Levin et al. 2007; Cachat et al. 2010; Gebauer et al. 2011) as a validated measure of anxiety-like behavior in adult zebrafish (**Fig. 3a**, see Methods for details). In the NTD task, WT fish explored the arena broadly (**Fig. 3b-d**). These findings parallel observations in the OF task and are consistent with a low-stress state and robust exploratory drive for WT fish.

When examining TALE and Hox mutant lines, the overall number of zone alternations did not differ significantly from WT fish (**Fig. 3b**). However, all TALE and Hox mutant lines spent significantly less time in the top, and more time in the bottom zone, than WT fish (p<0.05 or less for each genotype; **Fig. 3d**), indicating a shared elevation in anxiety-like behavior consistent with the OF findings. Accordingly, when examing the number of zone alternations specifically from the bottom to the top of the tank, Prep1 (p<0.01) and Hoxb1b (p<0.01) mutants made significantly fewer vertical transitions compared to WT (**Fig. 3c**), indicating impaired vertical exploration and increased thigmotaxic behavior similar to observations in OF.

Although shared elevation in anxiety-like behavior remained in all TALE and Hox mutant lines between the OF and NTD task, the nature of the stress response and coping strategies differed in some lines when compared across tasks. For example, in the NTD paradigm, Prep1 and TGIF1 mutants maintained the stress-responsive behaviors observed in the OF task, with Prep1 exhibiting significantly increased freezing (p<0.01) and TGIF1 displaying significantly elevated hyperactivity (p<0.0001, **Fig. 3e-f**). Interestingly, stress responses in some lines shifted in the NTD task compared to the OF. Hoxb1b mutants, which were hyperactive in OF, exhibited significantly increased freezing (p<0.0001) in NTD; **Fig. 3e-f**). The Pbx2, Pbx4, and Hoxb1a mutants exhibited hyperactivity in NTD, but not OF (**Fig. 3e-f**), consistent with a heightened active or hyper-aroused response only in the NTD task (Sumathipala et al. 2024). Taken together, these findings highlight that TALE and Hox genes not only modulate overall anxiety levels but may also influence the flexibility of stress coping strategies across environments and the way those environments are perceived as stressors.

### 3.3 Baseline and Stress-Evoked Social Behaviors Are Differentially Modulated by TALE and Hox Mutations

To assess baseline and stress-modulated sociability, we evaluated behaviors in groups of fish (**Fig. 4a**). Zebrafish are highly social animals that naturally form cohesive groups (shoals) whose structure is dynamically modulated by their physiological state and perceived threats (Speedie and Gerlai 2008). Such interactions are critical for survival and provide many benefits for normal life processes ((Magurran 1990; Griffiths and Magurran 1997; Killen et al. 2017) see Methods for details). When measuring sociability at baseline, groups of WT fish maintained consistent IID spacing (**Fig. 4b**), responding to each other’s movements while showing limited cohesion (**Supplemental Video 1**), suggesting that under non-stressful conditions, WT zebrafish engage in moderate shoaling (see Methods). Groups of Prep1 mutant fish exhibited a 32% increased IID (**Fig. 4b**, p<0.0001), and reduced group organization, cohesion, and polarization compared to WT (**Supplemental Video 1**). This diminished group coordination is unlikely to indicate a lower stress state, as zebrafish typically maintain baseline shoaling even under relaxed conditions (Buske and Gerlai 2011). Instead, it may reflect impaired social cue perception or disrupted neural integration of sensory information, resulting in reduced motivation or ability to engage in coordinated group movement. In contrast, groups of TGIF1 (p<0.0001) and Pbx2 (p<0.0001) mutant fish exhibited more coordinated behaviors at baseline (**Fig. 4b**), indicated by reduced IID and increased group organization and clustering at the bottom of the tank compared to WT (**Supplemental Video 2-3**), suggesting that TGIF1 and Pbx2 mutants display heightened cohesion and enhanced social awareness consistent with a stress-like behavioral state (Pitcher and Parrish 1993; Herbert-Read et al. 2011), even under baseline conditions. No differences relative to WT were observed for any other mutant lines (**Fig. 4b**).

Because certain mutant lines showed altered social behaviors at baseline, we next tested whether these differences persist or shift under stressful conditions. To address this, we employed the net chase paradigm ((Ramsay et al. 2009; Aponte and Petrunich-Rutherford 2019), see Methods for details). During net chase, groups of WT fish maintained, or slightly decreased, their IID (compare mean IID in **Fig. 4c** [7.84cm] to **Fig. 4b** [8.03cm]) along with becoming more polarized in motion, and aligning in direction, orientation, and speed (**Supplemental Video 4**). Groups of Prep1 mutant fish exhibited reduced coordination relative to WT during the net chase, similar to their reduced shoaling at baseline, as indicated by increased IID (p<0.0001; **Fig. 4c**), and decreased group organization, cohesion, and polarization (**Supplemental Video 4**). Groups of TGIF1 and Pbx2 (p<0.0001 for both) mutant fish exhibit enhanced cohesion compared to WT both at baseline and in the net chase paradigm (**Fig. 4b, c**). Lastly, Hoxb1b mutant fish showed a similar IID to WT at baseline, but had a significant (p<0.01) reduction in their IID relative to WT during the net chase (p<0.01). However, this effect reflected increased lethargy over the course of the net chase paradigm, resulting in all individuals settling to the bottom of the tanks, unmoving (**Supplemental Video 5**; see below), suggesting that additional physiological factors affecting aerobic capacity, energetic demand, or neuromuscular endurance outside of the stress response may be affected by these TF mutations,. No differences relative to WT were observed for any other mutant lines (**Fig. 4c**).

### 3.4 Stress-Evoked Lethargy in Select TALE and Hox Mutants Reveals Divergent Physiological Responses

In addition to the sociability changes observed during the net chase paradigm, several mutants also exhibited stress-induced lethargy that was not present at baseline. Fish that exhibited lethargic responses during the net chase task had reduced swimming activity and remained largely stationary, or moved only sporadically, either hovering near the bottom or floating passively along the top of the water column. These fish also had difficulties swimming against the current generated by the sweeping of the net, a loss of buoyancy control, abnormal postures, and labored breathing. We note that impaired group synchrony increases energetic costs and fatigue, potentially amplifying both behavioral and physiological dysregulation (Plaut 2001) and potentially contributing to the increased lethargy observed. At baseline, lethargy is not observed in any mutant lines (‘immobility’ category in **Fig. 5a**) with Prep1 mutants behaving like WT, Pbx4 and Hoxb1b mutants being slightly more active than WT, and Pbx2, Hoxb1a and TGIF1 mutants being considerably more active than WT (highly mobile category in **Fig. 5c**), consistent with the latter lines also displaying hyperactivity in other contexts (**Fig. 3f**). WT fish did not display lethargic behavior at any point during the net chase and returned rapidly to homeostatic conditions and normal shoaling behavior once the net chase ended (**Fig. 5b, d**; **Supplemental Video 6**). The Pbx4, and Hoxb1b mutant lines, that displayed modestly increased activity at baseline, exhibited increased lethargy relative to WT during net chase (p<0.0001 for all, **Fig. 5b, d; Supplemental Video 6**), while Pbx2, Hoxb1a and TGIF1 displayed similar heightened activity during net chase as observed at baseline (**Fig. 5a-b**). We conclude that, in addition to differential roles for TALE and Hox TFs in coordinating social behaviors, several of these TFs are also involved in regulating stress-induced lethargy.

**Fig. 5.**
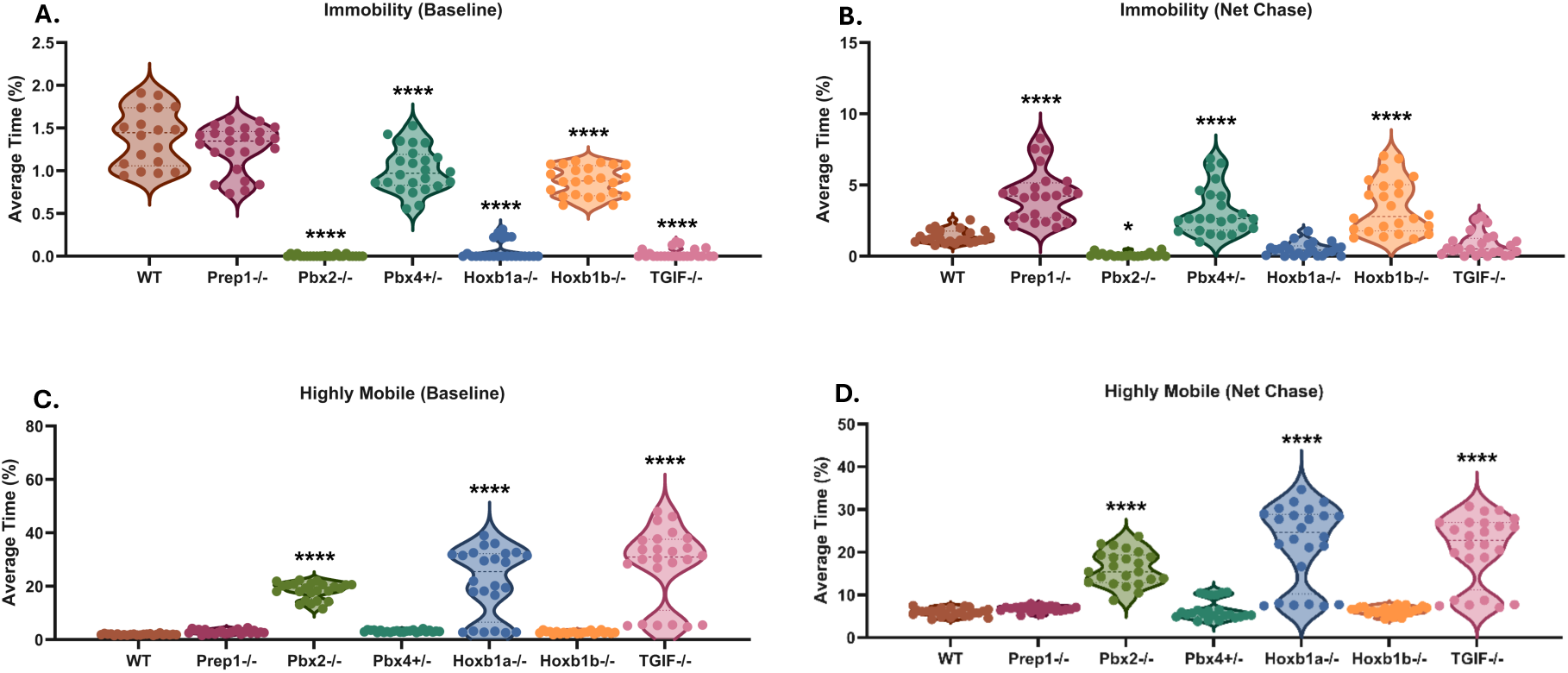
The Net Chase Paradigm Reveals Contrasting Physiological Responses in TALE and Hox Mutants. Violin plots showing percent time, averaged across the 10min time period, spent immobile (lethargy) at a) baseline and b) net chase, and percent time, averaged across the 10min time period, spent hyperactive at c) baseline and d) net chase. Each data point represents one animal. Data were analyzed using one-way ANOVAs. Significant differences compared to WT: *p<0.05, ****p<0.0001

### 3.5 Sex and Zygosity Do Not Influence Behavioral Disruptions in TALE and Hox Mutants

For all behavioral tests in this study, sex was considered in the analyses, but no sex-specific effects were found within each line (**Supplemental Fig. 2a-b**). Additionally, we examined potential differences in heterozygous versus homozygous mutations for each TALE and Hox line (aside from Pbx4, as noted above). We found no zygotic-specific differences within each line as the same disordered behaviors were observed in heterozygous and homozygous mutants within each line (**Supplemental Fig. 2a-b**). Therefore, the behavioral phenotypes observed are robust and independent of sex and a partial disruption of these developmental TFs is sufficient to produce measurable behavioral alterations.

### 3.6 Summary

Our findings reveal that mutations in TALE and Hox genes differentially disrupt key behaviors relevant to mental health disorders, including memory, stress responsivity/coping, and social behavior. Hoxb1a and Hoxb1b mutants exhibited impaired object recognition memory, suggesting deficits in cognitive processing or novelty detection.

Across stress-related assays, all mutant lines showed heightened anxiety-like behavior; however, distinct coping strategies emerged, with some lines favoring passive (e.g., Prep1) or hyperactive (e.g., TGIF1) responses.

Interestingly, stress-induced coping strategies in some lines shifted across tasks, suggesting task-dependent shifts in the stress-response strategy or stress perception. Sociability assays further revealed mutation-specific effects on baseline and stress-evoked group behavior, with Prep1 mutants showing reduced group cohesion and TGIF1 and Pbx2 mutants demonstrating enhanced group cohesion. Notably, these behavioral disruptions were independent of sex and were present in both heterozygous and homozygous states. It is important to consider that differences in locomotor activity, sensory perception, stress sensitivity, or motor capacity may contribute to these phenotypes, and additional assays will be necessary to disentangle primary effects on social behavior or cognition from secondary consequences of disruptions to other processes. Overall, these results highlight divergent and shared behavioral phenotypes resulting from TALE and Hox gene disruption (**Fig. 6**), offering insight into the gene-specific regulation of neurobehavioral function.

**Fig. 6.**
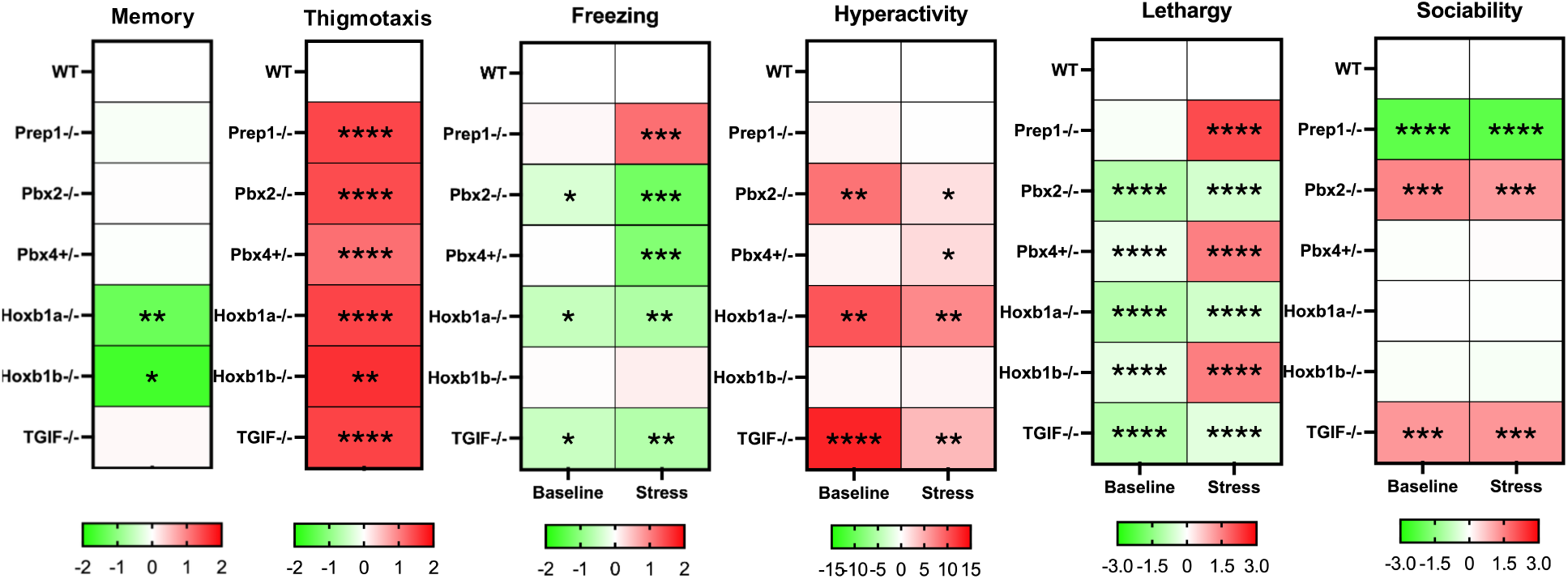
Comparison of Unique and Shared Behavioral Phenotypes Across Mutant Lines Heatmap illustrating behavioral metrics measured across each paradigm for each mutant line compared to WT. Data were analyzed using one-way ANOVAs. Significant difference compared to WT: *p<0.05, **p<0.01, ***p<0.001, p<0.0001

## 4. Discussion

Mental health disorders in humans rarely display isolated symptoms; instead, they emerge from complex interactions between genetic, developmental, and environmental factors. This can produce multi-domain impairments including, but not limited to, cognition, emotional regulation, stress responsivity, and social behavior. Our findings demonstrate that mutations in TALE and Hox genes—critical transcriptional regulators of early brain patterning (Mann and Affolter 1998; Waskiewicz et al. 2001, 2002; Choe et al. 2002; Gongal and Waskiewicz 2008; Ladam et al. 2018)—yield diverse but consistent behavioral disruptions in zebrafish, a model with conserved vertebrate neural circuitry. We show that mutations in TALE and Hox genes produce both shared and divergent disruptions across recognition memory, stress responsivity, and social cohesion in zebrafish (**Fig. 6**). These behavioral alterations—spanning cognitive inflexibility, heightened anxiety-like states, divergent coping strategies, and altered sociability—mirror the multidimensional impairments observed in human mental health disorders (Chand and Marwaha 2025; Hany and Rizvi 2025; Hodis et al. 2025; Musullulu 2025). Importantly, these phenotypes emerged largely independent of sex and were evident in both heterozygous and homozygous states, suggesting that partial disruption of these developmental TFs is sufficient to produce measurable behavioral alterations, potentially through changes in neural development or circuit function. This is consistent with observations in humans that many genetic variants underlying neuropsychiatric risk are haploinsufficient, with a single defective allele being enough to precipitate functional deficits (Shibutani et al. 2017; Earl et al. 2017; van der Sluijs et al. 2019). Past studies in mice further underscore this principle, as Meis1 mutation alters motor phenotypes (Salminen et al. 2017; Kittke et al. 2024) and nociceptor development (Cao et al. 2022), highlighting a role for TALE-family TFs in regulating both physiological processes and complex behaviors.

The zebrafish phenotypes observed here align with comorbidity patterns frequently seen in human neurodevelopmental and neuropsychiatric disorders, where impairments rarely occur in isolation but instead span multiple behavioral domains (Sanders et al. 2019; Farooqui 2021; Antolini and Colizzi 2023). Hoxb1a and Hoxb1b mutants demonstrated impaired recognition memory, evidenced by reduced exploration of the novel object. This shared phenotype may stem from disruptions to conserved functions for Hoxb1a and Hoxb1b in hindbrain patterning (McClintock et al. 2002; Weicksel et al. 2014), which may have a broader impact on downstream circuits related to learning and memory, such as the dorsolateral pallium, the hippocampal-analogous circuitry in teleost fish, including zebrafish (Broglio et al. 2010). In mammals, reduced novelty preference is linked to hippocampal–prefrontal network dysfunction (Alemany-González et al. 2020), a hallmark of schizophrenia, autism spectrum disorder (ASD), and certain intellectual disabilities (Oh et al. 2017; Desaunay et al. 2023). Clinically, such deficits manifest as reduced cognitive flexibility, working memory impairments, and difficulty integrating new information—features that can exacerbate emotional dysregulation and stress sensitivity (Raio et al. 2017; Peckham et al. 2019; Groves et al. 2020). Interestingly, while Hoxb1a and Hoxb1b mutants both displayed deficits in recognition memory, they diverged in their stress-response profiles. This divergence may reflect differences in transcriptional partnerships and genomic targets despite their close paralogy. Thus, phenotypic overlap in one behavioral domain (cognition) does not necessarily predict convergence in another (stress).

Across mild stressor tasks (OF and NTD), all mutant lines displayed increased thigmotaxis, indicating a shared elevation in anxiety-like behavior, and suggesting that TALE and Hox genes may collectively support development of neural circuits underlying basal stress responsivity. Yet, their coping strategies in response to stress diverged by genotype. Prep1 mutants exhibited passive coping characterized by freezing, immobility, and reduced vertical exploration—behaviors reminiscent of depressive withdrawal, PTSD-related freezing, or behavioral inhibition in humans (Schmidt et al. 2008; Volchan et al. 2017). In contrast, TGIF1, Pbx2, and Hoxb1a mutants adopted active or hyper-aroused coping, paralleling hypervigilance, agitation, or maladaptive escape behaviors seen in generalized anxiety disorder or hyperarousal states in PTSD (Cantor 2009; Mahoney et al. 2018). Certain mutants, such as Hoxb1b and Pbx4, transitioned between hyperactive and passive strategies depending on the context. This behavioral variability may reflect underlying differences in neural circuitry engagement, with certain mutants showing rigid or maladaptive responses to novel contexts. These coping style differences may arise from TF-specific influences on stress-regulatory systems.The hypothalamic–pituitary–interrenal (HPI) axis (the zebrafish equivalent of the mammalian HPA axis (Lee et al. 2024)) together with limbic and monoaminergic circuitry, represent conserved vertebrate systems known to regulate responses to stress (Stephenson-Jones et al. 2012; Ortega et al. 2021). Whether these pathways are disrupted in TALE and Hox mutants remains an important direction for future investigation. Thus, even where multiple TFs converge at the level of anxiety-like behavior, they may diverge in their roles within circuitry that regulates active versus passive stress responses.

Social behavior assays revealed further bidirectional effects. Prep1 mutants showed reduced group cohesion at baseline and under stress, consistent with social withdrawal, diminished responsiveness to group cues, or impaired integration of social signals—paralleling ASD and the negative symptom cluster of schizophrenia (Yamada et al. 2023; Fulford and Holt 2023). In contrast, TGIF and Pbx2 mutants exhibited heightened group cohesion and organization, potentially reflecting increased affiliative drive or over-synchronization of social behavior, features observed in certain ASD subtypes with rigid adherence to social norms (Key et al. 2022; Petrolini et al. 2023). Stress-induced lethargy was observed in Hoxb1b, Prep1, and Pbx4 mutants during the net chase, with fish displaying waning swimming capacity, becoming immobile, losing buoyancy control, and having abnormal postures. Further studies are needed to fully characterize these physiological stress responses and identify the additional systems that may be involved. Conversely, hyperactive genotypes (e.g., TGIF1 mutants) sustained elevated activity throughout stress exposure. Therefore, future investigations into how TF disruption can influence physiological components of the stress response, such as motor, neuromuscular, or metabolic systems should be explored.

Taken together, these findings reveal both overlapping and non-redundant roles of TALE and Hox TFs in shaping vertebrate behavior (**Fig. 6**). Shared phenotypes, such as elevated anxiety-like responses, likely arise from common developmental functions and cofactors within the TALE and Hox families. In contrast, mutation-specific behavioral profiles, such as cognitive inflexibility, altered stress coping, and changes in social interaction, may reflect the molecular specialization of individual TFs, including unique genomic targets, expression patterns, and protein complex formation. For example, Hoxb1a and Hoxb1b are highly related based on sequence identity and share some functions (novel object exploration), but differ in others (stress coping responses). Our findings therefore suggest that TF family members are unlikely to be fully interchangeable in establishing functional behavioral neural circuitry. Consequently, TF family members may only partially compensate for one another—for example, Hoxb1a may substitute for Hoxb1b at certain genomic sites, but recruit different transcriptional complex components, which leads to misregulation of Hoxb1b targets and also reduces Hoxb1a availability at its normal binding sites. Thus, behavioral phenotypes may be shaped not only by the loss of a given TF, but also by inappropriate activities of related TFs and their complexes. Ultimately, this framework clarifies how subtle differences in TF family function may shape divergent neurobehavioral outcomes, advancing our understanding of genetic contributions to mental health vulnerability.

## Notes

**Funding** This work was supported by the National Institutes of Health grant R01GM142158 to CGS and grant R35GM159847 to JCN.

### Competing Interest Statement

The authors have declared no competing interest.

