## Supplemental Figures for "TALE and Hox Transcription Factors Control Adult Behaviors in Zebrafish"

### Files included:

### Supplemental Table 1

### Supplemental Figures 1 and 2

### Supplemental Videos 1 - 6

**Supplemental Table 1: Characteristics of TALE germ line mutant alleles.** [a]Nucleotide sequence change of mutant alleles. Deletions are shown as dashes, insertions as bold, and point mutation changes in red. Wild type (top) and mutant (bottom) sequences are shown for each mutant. [b]Amino acid sequence change shown in red (premature stop codon shown with an ⊗). [c]Indicates whether mutation results from insertion, deletion, or point mutation. Numbers in parenthesis indicate net gain/loss of nucleotides in mutant sequence. [d]Guide RNA sequences used for Cas9 mutagenesis. [e]Method used to induce mutation.

| Gene | Nucleotide Sequence[a] | Amino Acid Sequence[b] | Type of Mutation[c] | Guide RNA Sequence[d] | Method Type[e] |
|---|---|---|---|---|---|
| *prep1.1*<br>*(co77)* | GTCACACCACAGGGACAAGTGGTGACACAGGCCATCTCAC<br>GTCACACCA-------------------------TCTCAC | VYQPVTVVTPQGQVVTQAISPGALHIQNSQL<br>VYQPVTVVTPSHLEHYTSRTHSSNFSSIRT⊗248 | Deletion<br>(-25bp) | TCACACCACAGGGACAAG | CRISPR/Cas9 |
| *pbx2*<br>*(co126)* | AGCCGGGCCAGAGA AAGGGGGCGGAGC<br>AGCCGGGCCAGAGA**GGGAAGGGGGCGG**AAGGGGGCGGAGC | GVAGPEKGGGAAAAVSAATSSGGMSPDSSLE<br>GVAGPEREGGGRGRSRSGSICCDQLWRDVP⊗165 | Insertion<br>(+13bp) | TGTAGCCGGGCCAGAGAAAG | CRISPR/Cas9 |
| *pbx4*<br>*(co127)* | GGCGTCTCTGGGCCAG AGAA GGGTGGAGGATCT<br>GGCGTCTCTGGGCCAG**GT**AGAA**AATCT**GGGTGGAGGATCT | LAEGVSGPEKGGGSAAAAAAAAAAGGSPNDG<br>LAEGVSGPGRKSGWRICGGSSSCSGSWRIT⊗141 | Insertion<br>(+7bp) | GGCGTCTCTGGGCCAGAGAA | CRISPR/Cas9 |
| *TGIF1*<br>*(fh258)* | AAGACCGCAACAGCTACGAGCCTGGCTCACCTCACCCGAC<br>AAGACCGCAACAGCTAAGAGCCTGGCTCACCTCACCCGAC | RRGSKGGEMLSDNSQSPKHGLLANGEDRNSY<br>RRGSKGGEMLSDNSQSPKHGLLANGEDRNS⊗143 | Point<br>Mutation | N/A | ENU |

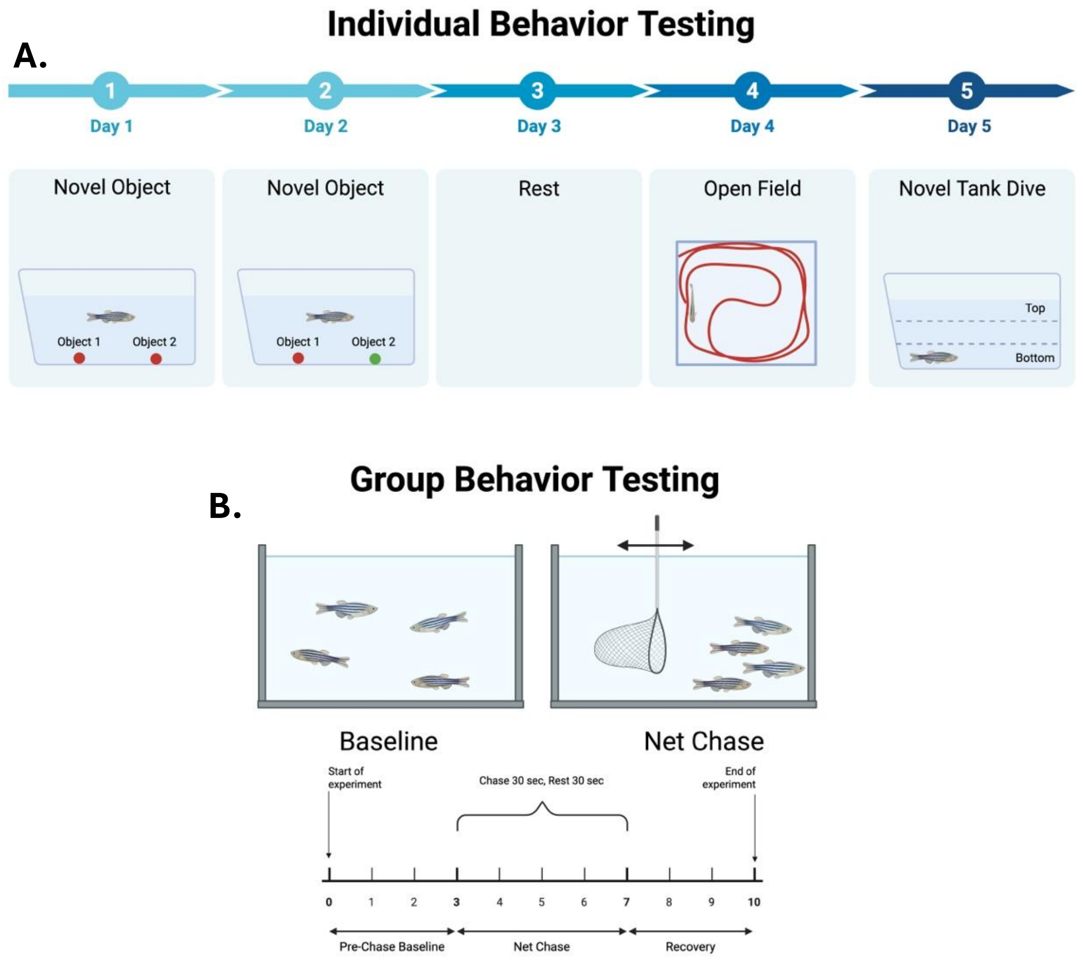


**Supplemental Figure 1: Experimental Timeline of Behavioral Testing.** Schematics illustrating the experimental timeline for **A)** Paradigms used for individual behavioral testing, and **B)** Paradigms used for group behavioral testing.

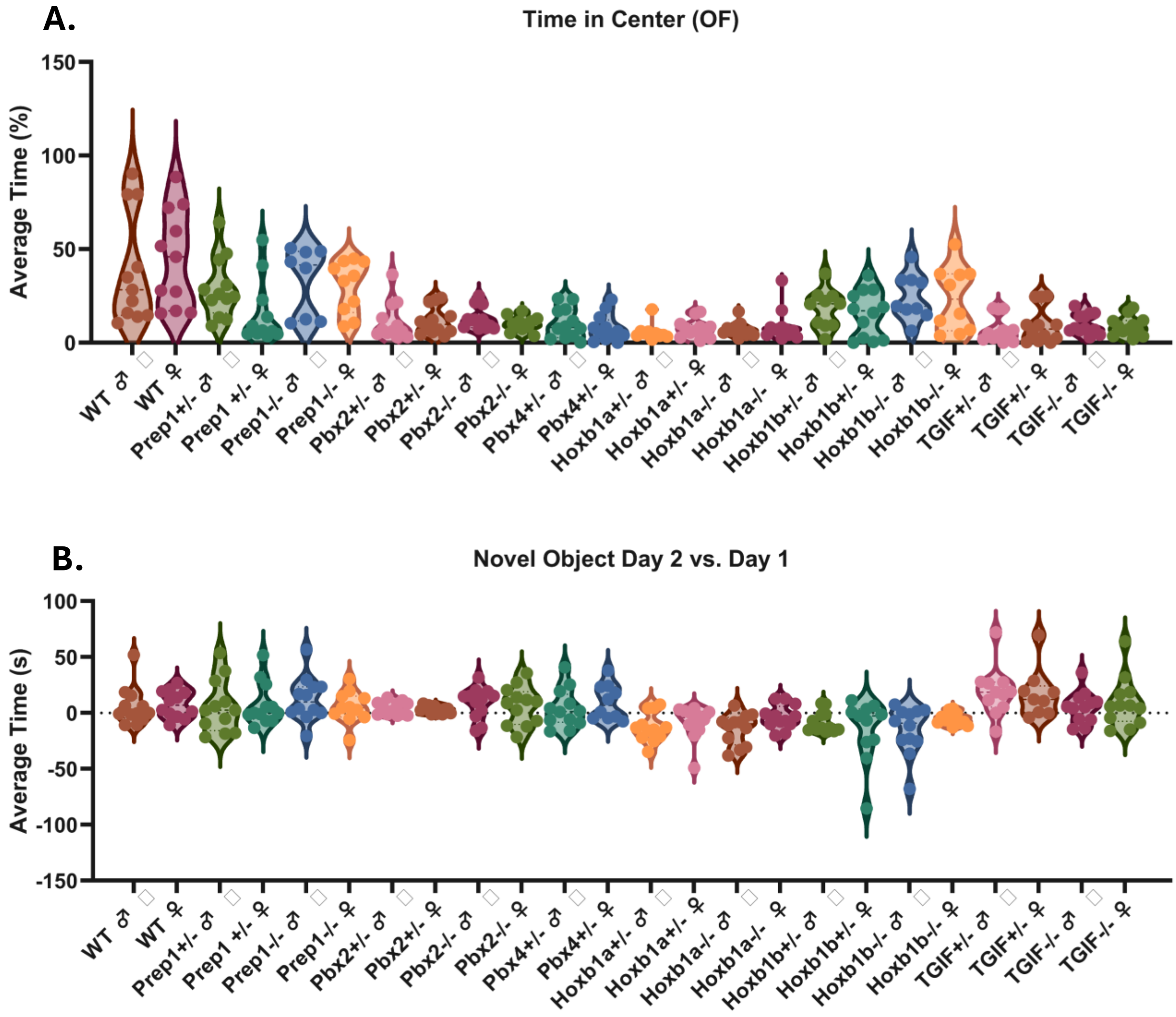


**Supplemental Figure 2: Sex and Zygosity Do Not Affect Behavior.** Bar graphs showing male and female performance within WT and TALE and Hox mutant heterozygous and homozygous adult zebrafish in **A)** Open Field (OF) and **B)** Novel Object Recognition Task (NOR), where a positive number equals more time around the novel object and a negative number is less time around the novel object. Comparisons were made between heterozygous and homozygous mutants (where applicable), and between male and female fish, for each line. Data were analyzed using two-way ANOVAs. No significant differences were found between males and females, or heterozygous and homozygous mutants within each line.

[Supplemental Video-1]

[Supplemental Video-2]

[Supplemental Video-3]

[Supplemental Video-4]

[Supplemental Video-5]

[Supplemental Video-6]

**Supplemental Videos 1-6: TALE and Hox Mutants Exhibit Altered Social Behaviors and Stress-Induced Lethargy Compared to Wild-Type Controls.** Representative videos exhibiting alterations in social and physiological responses between WT controls and TALE and Hox mutant lines during baseline and the net chase paradigm for **1)** Reduced shoaling behaviors in Prep1.1 mutants compared to WT controls at baseline, **2)** Enhanced schooling behaviors at baseline in TGIF1 mutants, **3)** Enhanced schooling behaviors at baseline in Pbx2 mutants, **4)** Reduced schooling behaviors in Prep1.1 mutans compared to WT controls during the net chase paradigm, **5)** Increased lethargy in Hoxb1b mutants during the net chase paradigm, and **6)** Examples of postures associated with lethargy in mutant fish compared to WT controls following the net chase paradigm.
